# Deep Learning-Augmented Stimulated Raman Imaging for Cell-Type-Specific Metabolic Profiling in Live Neuronal Co-Cultures

**DOI:** 10.1101/2025.11.18.689078

**Authors:** Li-En Lin, Xiaotian Bi, Adrian Colazo, Haomin Wang, Lu Wei

## Abstract

Neuronal metabolism is fundamental to brain functions and diseases, yet its spatial and temporal dynamics and interactions remain poorly understood. Here, we introduce a tandem deep-learning approach integrated with bioorthogonal chemical imaging using stimulated Raman scattering (SRS) microscopy. This method achieves high-speed and quantitative metabolic profiling in live neuronal co-cultures. Our deep-learning framework consists of a recurrent convolutional neural network (RCNN) that enables high-resolution 3D imaging with minimal photodamage and a U-Net segmentation model for cell-type-specific metabolic analysis. Using deuterium-labeled metabolites, we demonstrate the ability to trace lipid, protein, glucose, and D_2_O metabolism in neurons, astrocytes, and oligodendrocytes under physiological and pathological conditions, including NMDA receptor activation, proteasome inhibition, and Huntington’s disease. Our findings reveal distinct metabolic adaptations among neuronal cell types and underscore the importance of non-invasive metabolic profiling for understanding neuronal interactions and disease mechanisms. This platform significantly advances live-cell dynamic imaging with broad applications in neuroscience, disease modeling, and therapeutic screening.

## Introduction

Neuronal metabolism plays a critical role in brain function, aging, and neurological disorders. Extensive evidence has suggested that metabolic activities, including lipid metabolism, energy (or glucose) metabolism, and protein metabolism, in the central nervous system are tightly regulated across spatial and temporal scales. Disruptions in these metabolic processes have been implicated in neurological diseases, such as autism, neurodegenerative diseases, and brain cancer^1–7^. However, metabolic processes involving intricate molecular types, spatial distribution, and temporal dynamics remain poorly characterized at the subcellular level across different neuronal cell types in their native states^8,9^. This knowledge gap hence hinders our understanding of normal brain functions, such as learning and memory, and pathological conditions.

Existing methods that probe neuronal metabolic activities face fundamental limits. Conventional MRI and PET lack the fine spatial resolution to resolve cellular structures. Mass spectrometry-based methods offer fine spatial resolution and high multiplex imaging (measuring tens of molecules simultaneously), but are invasive and incompatible with live-cell studies^10,11^. Fluorescence microscopy has live-cell biocompatibility with high spatial and temporal resolution. However, it commonly requires bulky fluorophore labeling, which introduces significant physical and chemical perturbations to the native function of much smaller metabolites. Genetically encoded fluorescent biosensors sensitively detect subcellular metabolic activities^12–15^. However, they are limited to a small number of metabolites, and permit observations only in transfected neurons or neuronal compartments, missing the global information^16–18^. Therefore, there is an urgent need for noninvasive, versatile, and quantitative imaging techniques capable of probing metabolic activities in the complex nervous system, providing multi-dimensional information under both physiological and pathological conditions.

A recent bioorthogonal chemical imaging platform has been introduced that couples SRS microscopy with small bioorthogonal vibrational tags^19^. This approach is uniquely suited for live neuron and brain tissue imaging with minimal perturbation to endogenous metabolic activities. For instance, it involves selectively replacing stable carbon-hydrogen (CH) bonds with carbon-deuterium (CD) bonds derived from small metabolites. SRS imaging then targets the specific vibrational frequency of CD bonds within the cell-silent region, which correspondingly reports distribution of either the original metabolites or the downstream metabolized biomass. As a result, this method has been established as a quantitative imaging platform for monitoring the uptake, synthesis, and turnover dynamics of key cellular metabolites, including amino acids, fatty acids, glucose, and small molecule drugs. (**Fig. 1a**)^20–24^. In addition to using biorthogonal labels, the label-free SRS targeting the endogenous CH_3_ (2940 cm^-1^) and the CH_2_ (2845 cm^-1^) vibrational channels have been well-established to ideally outline the morphology and capture the distribution of pre-existing total proteins and lipids in live neurons with intrinsic 3D sectioning^25,26^.

**Figure 1.**
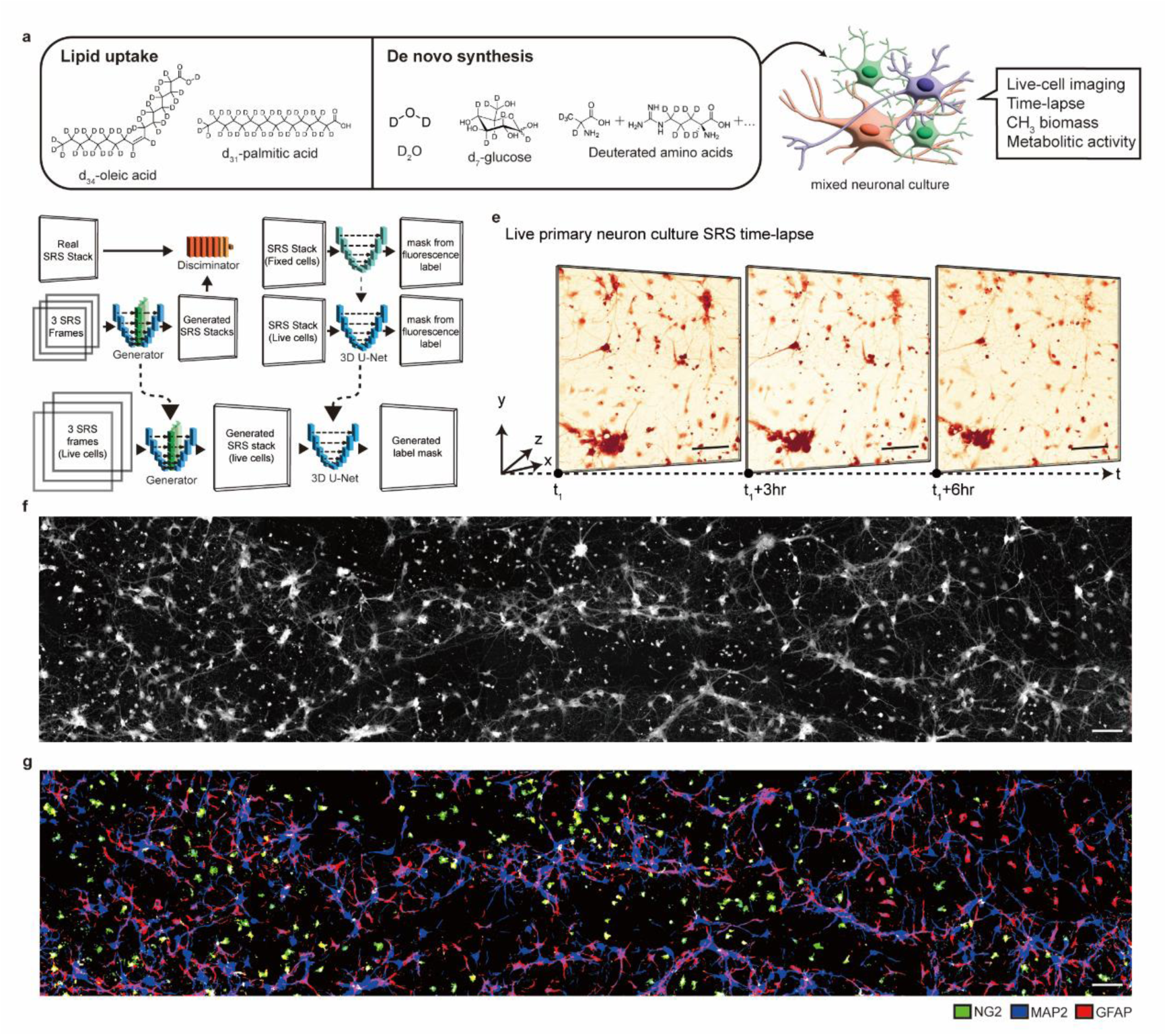
Experimental workflow. **(a)** Left: Structures of representative deuterated metabolites coupled with SRS imaging for assessing saturated and unsaturated fatty acid uptake and turnover, D₂O metabolic incorporation, glucose metabolism, and protein synthesis and turnover. Right: Schematic illustration of mixed neuronal co-cultures with intertwined morphology and spatial localization. **(b)** A scheme of the RCNN network for predicting the complete image stack based on preceding and succeeding frames. **(c)** A scheme of the U-Net model for segmenting images into different neuronal cell types. **(d)** An overall scheme for the implementation: in live-neuron experiments, three SRS frames were used to predict the entire stack using the model in **(b),** followed by segmentation utilizing the model in **(c)**. **(e)** Demonstration of robust live-neuronal time-lapse imaging with our optimized SRS acquisition workflow. **(f-g)** Maximum Z-projection of a large field-of-view mosaic stitch of the SRS images at CH_3_ (2940 cm^-1^) channel (f) of a live neuronal co-culture, with correspondingly predicted cell identities based on fluorescence-immuno-staining ground-truth of NG2 (neuron-glial antigen 2), MAP2 (microtubule-associated protein 2) and GFAP (glial fibrillary acidic protein) (g). Scale bars: 100 µm.

To probe neuronal metabolism, co-culture systems, such as neuron–astrocyte–oligodendrocyte cultures, offer high-resolution and well-controlled experimental conditions over animal models or brain tissues. It preserves the complex co-culture extracellular microenvironment. This allows for the investigation of distinct metabolic activities and facilitates comparison of metabolic heterogeneity across various cell types under identical conditions (**Fig. 1a**)^27, 28–31^. However, key limitations remain for bioorthogonal chemical imaging in neuronal co-cultures. First, it remains challenging to perform longitudinal SRS imaging for live and delicate neuronal systems, capturing whole 3D information without inducing excessive photodamage. Second, due to the diverse morphology of different neuronal cell types in a mixed culture, accurate identification of cell identities between individual cells in a non-invasive manner remains a non-trivial task^32,33^. In fluorescence, identifying neuronal identity commonly relies on immunostaining, which requires cell fixation. The metabolic impact of organic dye-based neuronal targeting remains unclear, and the approach may also suffer from non-specificity^34–37^. Addressing these two challenges is essential for realizing the full potential of quantitative live metabolic imaging across diverse neuronal cell types. It is also critical for evaluating hypotheses such as molecular transport across heterogeneous cell populations^38^. Current imaging strategies often rely only on 2D images, which sacrifice volumetric accuracy, or on dense volumetric imaging with fine z-steps, which imposes significant phototoxicity. Our approach provides a balance between these extremes by reconstructing 3D stacks from sparse input data, offering a sweet spot between resolution, acquisition time, and neuronal viability.

Here, we introduce a tandem deep learning strategy comprising two data processing model components, integrated with stimulated Raman scattering (SRS) imaging to address the above-mentioned fundamental challenges. This is followed by bioorthogonal SRS imaging for dynamic metabolic analysis (**Fig. 1b-d**). The first recurrent convolutional neural network (RCNN) component leverages the sequential and continuous nature of cells to predict fine 3D images using sparsely collected data^39^. This approach allows precise and longitudinal imaging of neuronal morphology in 3D, which is essential for informing the second component of the architecture. It also significantly reduces image acquisition time and minimizes photodamage to live neurons. The second component, a U-Net convolutional neural network (CNN) architecture, segments the volume and predicts corresponding cell-type-specific labels. Subsequently, biorthogonal chemical imaging and tracing facilitate quantitative analysis of targeted metabolic behaviors in individual live neuronal cells. This tandem approach enables non-destructive, real-time investigation of neuronal cell types within primary neuronal cultures. It also provides novel insights into live neuronal metabolism and dynamics across different cellular species within the same local environment. Our method should represent a significant advancement in the field of live cell dynamic imaging, offering a versatile tool for studying neuronal function and metabolism in a complex co-culture environment.

## Results

### High-speed live-cell image acquisition assisted by RCNN

The diverse morphology of neuronal cells leads to drastically different information across the z-dimension. For quantitative metabolic analysis, it is essential to capture the entire 3D volume to precisely determine the entire cell distribution. However, live neurons are especially susceptible to phototoxicity from prolonged light exposure, compared to bacteria or mammalian cells. Such consecutive z-scanning (up to 20 frames with 0.5 μm step size) needed for high-resolution SRS volumetric imaging with an axial resolution about 1 µm would compromise neuronal viability and prevent longitudinal imaging. Since this imaging process is a continuous temporal sequence, we reasoned it as a problem well-suited for recurrent neural networks (**Fig. 1b, d**)^39^ . Unlike traditional methods that either analyze only 2D slices or require complete acquisition of densely sampled z-stacks, our RCNN enables accurate volumetric reconstruction from sparsely sampled frames, effectively striking a balance between information completeness and imaging burden. By selectively skipping intermediate frames, an recurrent structure can be leveraged to predict the missing frames based on their preceding and succeeding frames. To enhance spatial accuracy in image reconstruction, convolutional components are integrated to refine the representation of cellular features. We anticipate that a substantial amount of cellular information can be effectively captured even from relatively sparse data, significantly reducing the burden of data acquisition without compromising essential biological insights.

We evaluated z frame intervals from 0.5 to 5 μm for label-free SRS imaging targeting the CH_3_ (2940 cm^-1^) channel. Our model training strategy utilizes image stacks sampled at consistent inter-frame intervals relative to the central. Due to the difficulty in consistently pinpoint exact location for optimal z position, these stacks are derived from axial (z) positions spanning –1 µm to +1 µm centered around the highest signal in the CH₃ channel (**Fig. S1**, left). A 4 μm interval was found to provide the most efficient and reliable predictions in neurons at developmental time points between *days in vitro* (DIV) 8 and DIV 12. Accordingly, we successfully trained our model to up-sample image stacks consisting of 3 frames acquired at 4 μm intervals into 17 frames at 0.5 μm intervals. We note here, unlike the static conditions of imaging fixed cells, live imaging involves continuous acquisition of dynamic cellular structures, where frame alignment and signal quality may vary due to real-time physiological fluctuations. This training approach significantly reduces acquisition time and enables the network to effectively interpolate intermediate frames and accommodate the variability inherent in live imaging inputs, thereby largely enhancing predictive accuracy (**Fig. S1**, right).

We compared our model with traditional interpolation methods and found that it consistently maintained lower prediction error across all frames, regardless of the variation in the starting frame of the input. Notably, it outperformed both spline and linear interpolation, particularly when the predicted frame was farthest from the ground truth (**Fig. S2**). Our implementation effectively accelerated the 3D imaging speed by 6-fold, which indeed enabled us to perform time-lapse imaging from the same batch of neuronal co-cultures with high cell viability (**Fig. 1e**). This approach offers a promising balance between imaging speed, data sparsity, and cell viability, paving the way for more efficient and less perturbative imaging techniques in longitudinal live-neuronal analysis (**Fig. 1f, Fig. S3a**).

### 3D Cell Segmentation in Neuronal Co-cultures

Following the establishment of high-speed 3D live SRS imaging, we next aim to segment the neuronal cell types to generate cell identity masks for metabolic analysis. We adopted a hippocampal neuronal co-culture model, with the specific goal of delineating neurons, astrocytes, and oligodendrocytes. For this segmentation task, we adopted U-Net, a highly effective model for medical image segmentation due to its ability to capture fine details and spatial hierarchies in biological images^40–42^. We trained the model with live-neuronal SRS images acquired at 2940 cm^-1^, with corresponding immuno-fluorescence images (e.g. Microtubule-associated protein 2 (MAP2) for neurons; Glial Fibrillary Acidic Protein (GFAP) for astrocytes; Neuron-glial antigen 2 (NG2) for oligodendrocyte precursor cells) after fixation as ground truth. In live-cell imaging, image stacks were acquired at 4 μm intervals and were up-sampled to an effective resolution of 0.5 μm using a RCNN before feeding the data into U-Net. This RCNN preprocessing step significantly improved our model’s ability to capture finer cellular morphology along the z-axis, essential for accurate 3D segmentation. (**Fig. 1g, Fig. S3b-3c**).

To validate the importance of our tandem RCNN-based resolution restoration and U-Net segmentation approach, we conducted a series of tests isolating each component. We trained U-Net using only the original 4-μm-spaced SRS stacks and the corresponding immuno-fluorescence images, bypassing RCNN, and compared it with models trained with full 0.5 μm stacks, and the model trained with RCNN-restored stacks. RCNN restored the predictive performance for neurons, astrocytes, and oligodendrocytes, compared to only using under-sampled data (**Table S1-S3**). These results highlight that with RCNN’s restoration of z-resolution, U-Net could more adequately capture the cellular structures needed for accurate segmentation without requiring additional images from the neuronal cells.

We next tested using three-frame SRS inputs to up-sample the z-resolution and directly predict fluorescence labels with the RCNN, bypassing the U-Net segmentation step. This approach again resulted in a noticeable reduction in segmentation fidelity. Specifically, when RCNN was directly trained to predict fluorescence labels, fractured pattern artifacts emerged, particularly in thinner structures such as astrocyte processes (**Fig. S4**). These findings suggest that decoupling interpolation from label prediction, as in our approach, facilitates more effective learning of structural patterns by the models. This tandem deep-learning approach hence empowers comprehensive 3D segmentation masks of unlabeled live neuronal co-cultures at rapid imaging speeds and with minimal photodamage.

### Quantification of Neuronal-Type Specific Metabolic Activity by Deep Learning-Augmented Deuterium-Labeled SRS metabolic Imaging

We then sought to apply the obtained segmentation masks from the model coupled with deuterium-labeled bioorthogonal SRS imaging for detailed quantitative metabolic profiling of each distinct neuronal cell type in live hippocampal co-cultures for the first time (**Fig. 1a**). After incubating the co-cultures in a custom medium containing deuterium-labeled metabolites for a defined period, we acquired images in the CH₃ channel for tandem model-based cell segmentation. Subsequently, we performed CD channel imaging to assess targeted metabolic activity across different cell types. In addition to the three neuronal markers described above, we further trained our model to predict seven distinct labels representing combinations of NG2, GFAP, and MAP2 expression. This was done to account for spatial overlap among neuronal cell types and to avoid underestimating the contribution of pixels that may correspond to more than one cell type. (**Figure S3&S5**).

To investigate the metabolic activities of each cell type, we incubated co-cultures with various deuterated metabolites, including deuterated- oleic acid (d-OA, 3 days), palmitic acid (d-PA, 3 days), amino acids (d-AA, 1 day), d-glucose (d-glucose, 5 days), and D_2_O (5 days), each targeting specific metabolic processes through tracking their incorporation and turnover, with incubation time optimized for CD signals. Here, d-OA and d-PA illuminate lipid metabolism, including uptake, storage, and breakdown, as these fatty acids are fundamental to lipid biosynthesis and membrane formation^43^. Additionally, d-AA provide insights into protein synthesis and degradation, revealing protein turnover, cellular stress responses, and metabolic adaptations^20,21^, and d-glucose here traces the metabolic fate of glucose into newly synthesized biomass through the tricarboxylic acid (TCA) cycle^22,44–46^. Finally, D_2_O serves as a broad metabolic marker for biomass through incorporation of D into the stable CD bonds, facilitating studies of proliferation, metabolic flux, and membrane dynamics^47–49^. SRS imaging of the corresponding CD frequency in each biomass species captures the localization and concentrations of these molecules for comparison among distinct neuronal types (**Fig. 2a-d**).

**Figure 2.**
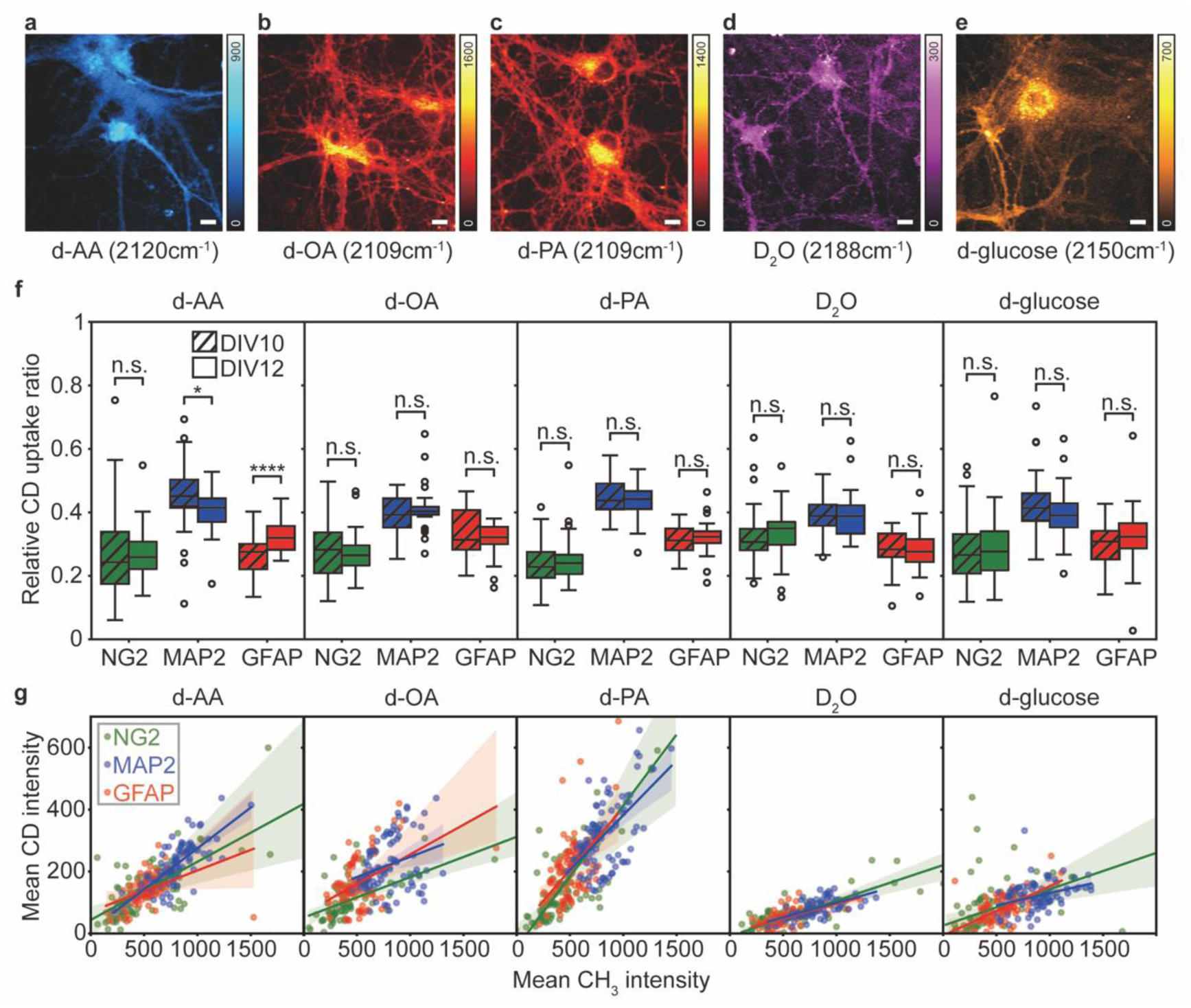
Quantification of metabolic incorporation across cell types with deuterated metabolite incubation. (a-e) Representative CD images of cells incubated with d-AA, d-OA, d-PA, D2O, and d-glucose. **(a)** Relative CD uptake ratio among predicted oligodendrocytes (NG2), neurons (MAP2), and astrocytes (GFAP) between imaging at DIV10 and DIV12 for d-AA (both 1-day incubation), d-OA, d-PA (both after 3-day incubation), D₂O, and d-glucose (both after 5-day incubation). The sum of CD ratios from all three cell types is equal to 1. **(b)** Observed positive correlations between metabolic incorporation (average CD intensity) and cellular protein density (average CH_3_ intensity) with different incorporation ratios from different metabolites. n.s., P > 0.05; *, P ≤ 0.05; ****, P ≤ 0.0001. Scale bars: 10 µm.

First, we performed metabolic activity analyses at both DIV10 and DIV12 using deuterated metabolites to compare across the three neuronal types. DIV10 and 12 are chosen since neuronal metabolism is expected to be active during this period while neuronal connections are already established *in vitro*. Interestingly, the relative CD uptake ratio across all three neuronal types (calculated by normalizing the sum of CD uptake ratios across all three cell types to 1), consistently indicated that neurons exhibited higher relative metabolic activity for all metabolites (**Fig. 2a** & **Fig. S6**, MAP2, blue), likely reflecting their higher metabolic demand in early development. Moreover, the relative CD uptake ratio further revealed a decrease in protein synthesis activity in neurons and a corresponding increase in astrocytes from DIV10 to DIV12, when considering the cellular microenvironment as a whole (i.e. 100% activity for all three neuronal types) (**Fig. 2a**, d-AA). In addition, although the relative lipid synthesis activity for d-OA does not have obvious change between DIV10 and DIV12 (**Fig. 2a**, d-OA), their absolute intensities indicate a consistent decrease of d-OA incorporation for all three neuronal types, indicating a decrease of total metabolic demands of OA for all neuronal cells at this developmental stage (**Fig. S7**).

Next, we analyzed the correlation between metabolic activity (i.e. CD intensity) and cellular protein density (i.e. CH_3_ intensity) across all three neuronal cell types. Interestingly, we observed positive correlations for all five metabolites. The difference in slopes indicates their relative turnover ratio (also indicating rates) when normalized to cellular protein levels (**Fig. 2b**). For instance, while both probing lipid turnover, d-PA presented a roughly two-fold steeper slope than that for d-OA (**Fig. 2b**, d-OA), possibly due to its greater involvement in membrane synthesis or storage lipid pools at the developmental stage. The slope of d-AA incorporation is comparable to that for d-OA. Again, neurons exhibit higher metabolic activities for protein synthesis among all three cell types. Although being incubated for five days prior imaging, both D_2_O and d-glucose exhibited limited incorporation with the lowest CD/CH ratios relative to all other probes (**Fig. 2b**, D2O and d-glucose). These observations from deuterated metabolite incorporation in neuronal co-cultures reveal distinct and heterogeneous patterns of metabolic activities. They emphasize the need for non-invasive metabolic profiling under controlled treatment and microenvironmental conditions.

### Neuronal Metabolic Activities with NMDA Receptor Activation and Excitotoxicity

NMDA receptors are pivotal in mediating excitatory neurotransmission and are critical for synaptic plasticity, memory formation, and learning^50^. These receptors mediate glutamate signaling by allowing calcium (Ca²⁺) and sodium (Na⁺) influx into neurons upon activation, thereby depolarizing the membrane and ultimately triggering the generation of action potentials^51^. The activation of NMDA receptors has been shown to be tightly coupled with increased neuronal activity, which imposes a substantial metabolic demand^52^.

Given that neurons predominantly rely on glucose as their principal energy substrate, we expect that this heightened activity drives an increase in glucose metabolism. We performed the metabolic analysis for all above selected five deuterated metabolites with and without 10 µM NMDA treatment. Our observations indeed indicate an increased uptake of d-glucose specifically in neurons, accompanied by elevated D₂O incorporation, reflecting an upregulation of selected metabolic processes in response to prolonged NMDA receptor activation (**Fig. 3a**, MAP2, blue). These findings support the hypothesis of enhanced metabolic activity under excitatory stimulation^53^. For astrocytes, although they also showed slightly elevated D₂O incorporation, their d-PA metabolism was slightly decreased, which implies that they may have shifted from uptake to *de novo* lipid synthesis (**Fig. 3a**, GFAP, red).

**Figure 3.**
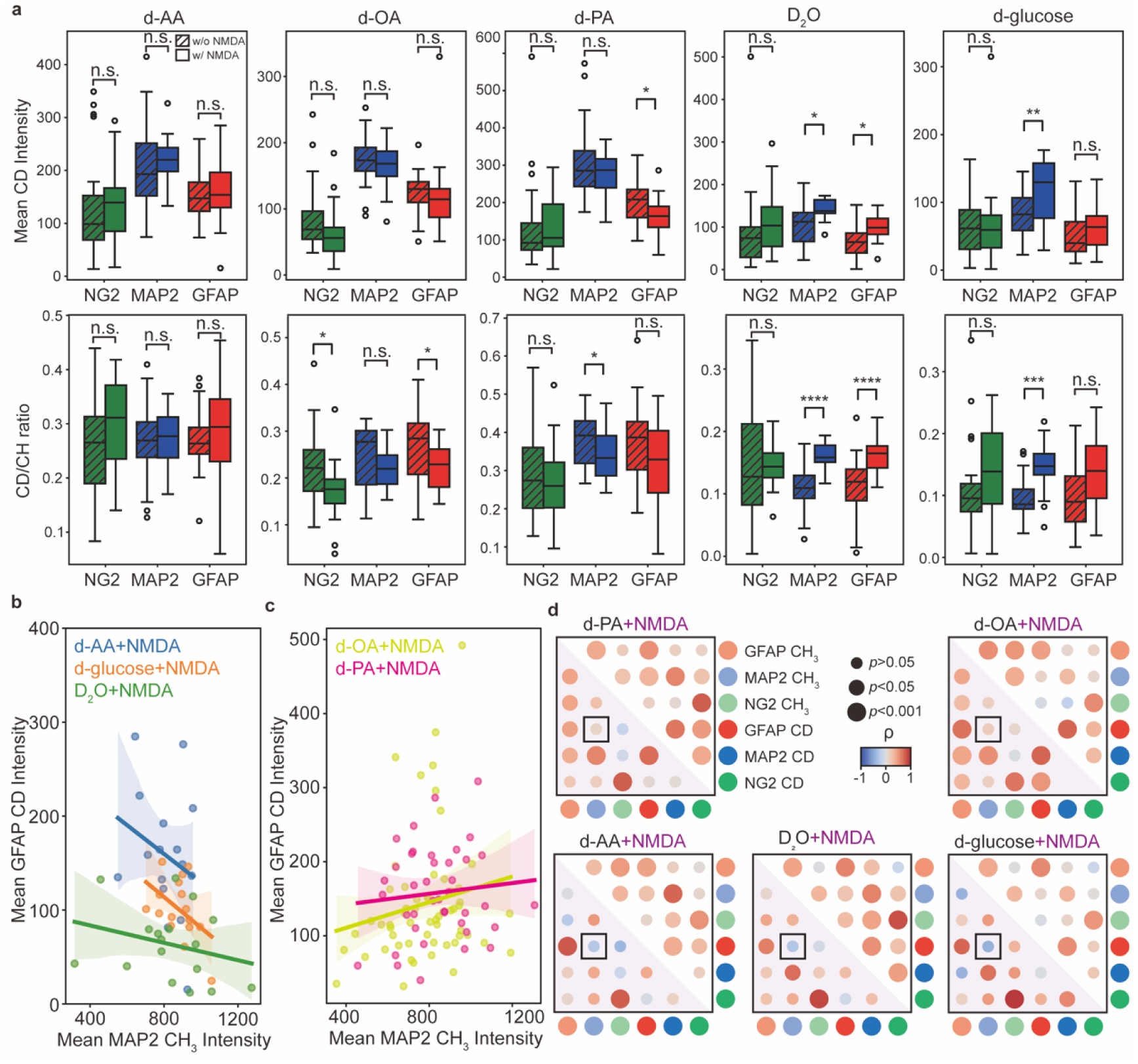
Effect of NMDA treatment on metabolic incorporation. **(a)** Comparison of mean CD intensity (top), and the CD/CH ratio (bottom) across three neuronal types in co-cultures incubated with (w/) and without (w/o) 10 µM NMDA treatment. **(b)** Negative correlation between GFAP-predicted CD intensity for *de novo* synthesis-related metabolites (d-AA, d-glucose, and D₂O) and MAP2-predicted CH₃ intensity. **(c)** Positive correlation between GFAP-predicted CD intensity for lipid species (d-OA and d-PA) and MAP2-predicted CH₃ intensity. **(d)** Correlation heatmap between metabolic incorporation (CD) and cellular protein density (CH_3_) in control neuronal cultures (upper right triangle) and NMDA-treated cultures (lower right triangle, pink shaded). Negative values (blue hues) indicate an inverse correlation, positive values (red hues) indicate a direct correlation. Black-boxed data presented in **(b, c)**. n.s., P > 0.05; *, P ≤ 0.05; **, P ≤ 0.01; ***, P ≤ 0.001; ****, P ≤ 0.0001. Scale bars: 10 µm.

Interestingly, we further observed a weak but consistent inverse relationship between neuronal bio-density (i.e. CH_3_ intensity from MAP2-predicted neurons) and astrocytic uptake of d-AA, D₂O, and d-glucose (**Fig. 3b, d**, black-boxed. In contrast, astrocytic metabolism of d-OA and d-PA showed a positive correlation with neuronal bio-density (**Fig. 3c, d**, black-boxed). Given the hypothesis that astrocytes support neuronal metabolism under stress conditions, this pattern suggests that astrocytes may adapt their nutrient utilization when surrounded by higher neuronal densities or when exposed to an extracellular environment with elevated neuronal protein content, as observed during NMDA-induced excitatory stress resulting from NMDA receptor overstimulation^54^.

### Cellular and Metabolic Responses to Proteasome Inhibition in Neuronal Co-cultures

Next, we investigated the impact of proteasome inhibition with 10 µM MG132 to metabolic changes. Inhibition of proteasome function results in a coordinated decrease in protein synthesis, mediated by a global proteostatic response in neurons^55^. In d-AA probed protein synthesis assays with MG132 treatment, we observed that the CD/CH ratio indicates a significantly reduced protein turnover rate (**Fig. 4a & 4b**, MAP2, blue). Surprisingly, the total CD intensity remains almost unchanged between treated and untreated neuronal cells (**Fig. 4a & 4b**, MAP2, blue). We found that all three neuronal cell types achieved the maintenance of total CD intensity through the reduction in cellular volume (**Fig. 4a**), likely reflecting their efforts to maintain homeostasis under MG132-induced stress through concentrating the newly synthesized proteins (**Fig. 4b**, d-AA).

**Figure 4.**
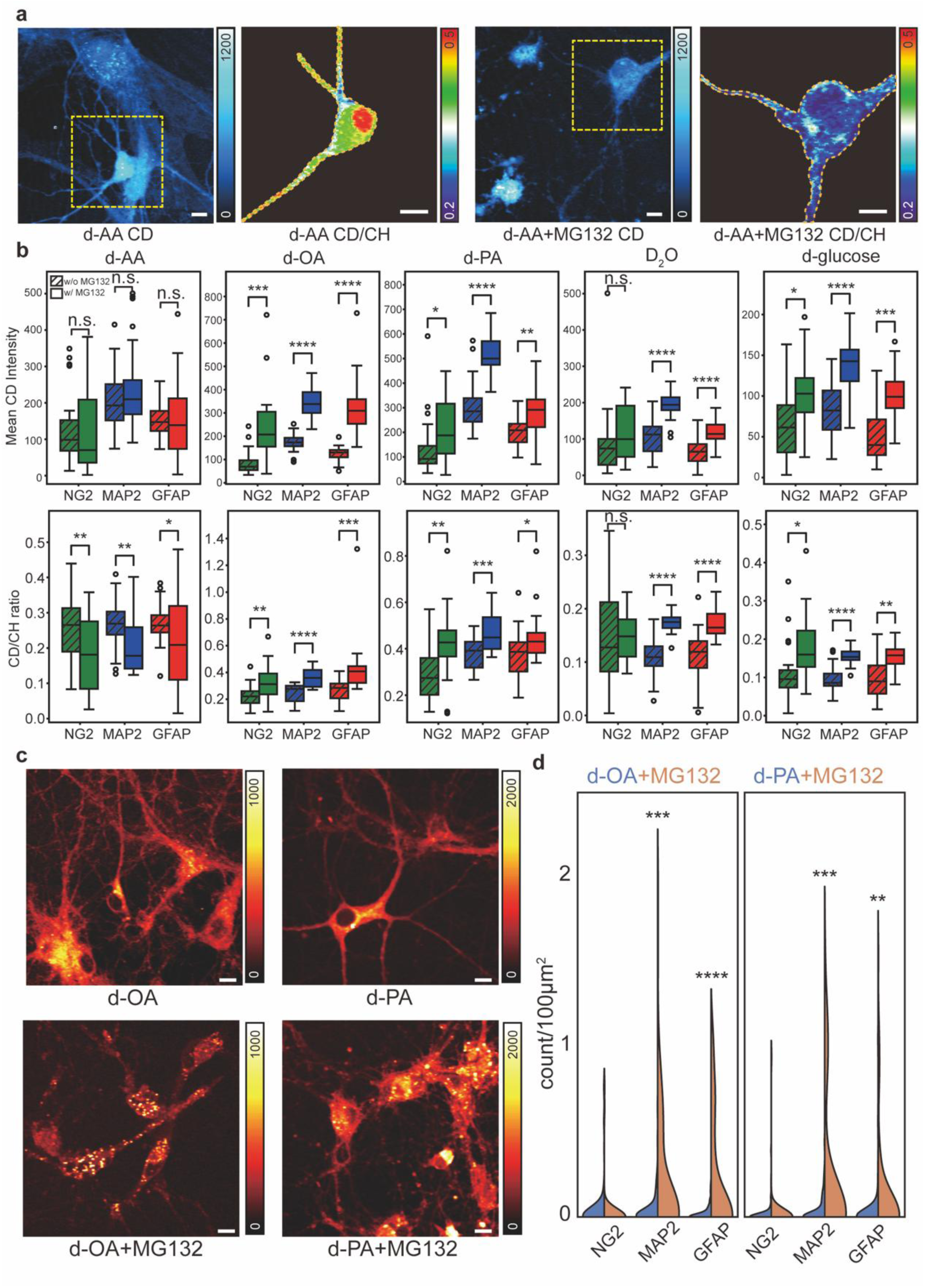
Effect of MG132 for proteasomal inhibition on neuronal metabolism. **(a)** Representative CD and CD/CH images of neuronal co-cultures probed with d-AA and co-incubated without and with 10 µM MG132. CD/CH images correspond to the boxed regions in the CD images. **(b)** Comparison of mean CD intensity and the CD/CH ratio between co-cultures incubated with and without 10µM MG132 treatment. **(c)** Representative CD images of co-cultures incubated with d-PA or d-OA, with or without 10 µM MG132. Increased lipid droplet accumulation was observed in the MG132-treated group (bottom), as quantified in **(d)**. **(d)** Violin plot of lipid droplet count per area for NG2, GFAP, and MAP2 predicted cells in control (blue) and MG132-treated (orange) groups. n.s., P > 0.05; *, P ≤ 0.05; **, P ≤ 0.01; ***, P ≤ 0.001; ****, P ≤ 0.0001. Scale bars: 10 µm.

Contrary to expectations, further metabolic analysis revealed increased metabolic incorporation of all four other metabolite species across all three neuronal types (**Fig. 4b**), likely function as a metabolic compensation under the proteasome stress. Notably, both d-PA and d-OA showed drastically elevated incorporation rate under MG132 treatment (**Fig. 4b**, 1.5- to 2-fold signal increase in d-PA and 2- to 3-fold for d-OA). These observations aligned with the significantly increased formation of lipid droplets (**Fig. 4 c-d**), a well-documented response to cellular stress and toxicity. The higher demand for oleic acids in response to MG132 stress also aligns with the neuroprotective properties of oleic acid for preserving cellular integrity. All these neuronal metabolic observations again confirm the necessity and the robust analysis of our approach.

### Metabolic and Cellular Stress Responses in a Neuronal Neurodegeneration Model of Huntington’s Disease (HD)

Finally, we applied our platform to study the cellular metabolic behavior of neuronal cells in a neurodegeneration disease model. We utilized adeno-associated virus (AAV) vectors to transduce neuronal co-cultures to produce mutant-Huntingtin (mHtt)-97Q-EGFP protein, a model construct commonly used by fluorescence imaging to study polyglutamine (polyQ) protein aggregation implicated in HD. Fluorescence microscopy was employed to identify cells forming protein aggregates with EGFP, and live neurons were subsequently imaged by SRS for segmentation and probing of the metabolic activities (**Fig. 5a**).

**Figure 5.**
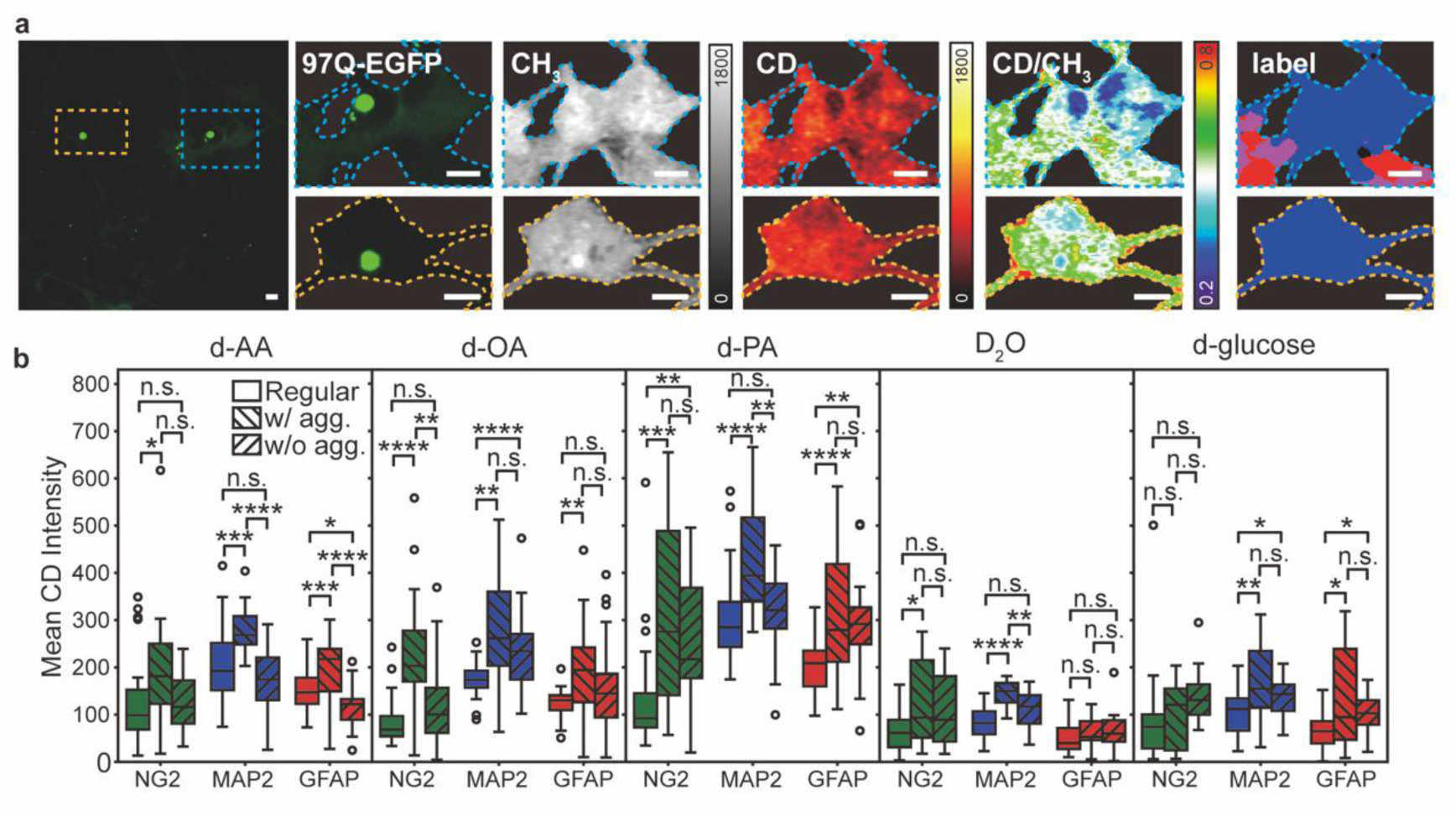
Impact of polyQ aggregation on neuronal metabolic incorporation. **(a)** Representative fluorescence, CH_3_, CD, CD/CH images, with predicted label images (red: GFAP, astrocyte; blue: MAP2, neuron) of neuronal co-cultures presented with polyQ aggregates while incubated with d-OA. **(b)** Comparison of mean CD intensity across three groups: non-transduced cultures (Regular), AAV-transduced cultures with protein aggregation (w/ agg.), and AAV-transduced neuronal cells without protein aggregation (w/o agg.). n.s., P > 0.05; *, P ≤ 0.05; **, P ≤ 0.01; ***, P ≤ 0.001; ****, P ≤ 0.0001. Scale bars: 10 µm.

Our results reveal that mHtt-97Q protein aggregation induces metabolic reprogramming (**Fig. 5b and Fig. S8**). Most obviously, elevated d-PA incorporation is shown for neuronal cells formed with polyQ aggregates (**Fig. 5b**). It is possible such increased metabolic demand is needed to support membrane repair that was disrupted by polyQ peptide^56^. Oleic acid levels were found to increase across all cell types (**Fig. 5b**), especially in oligodendrocytes, likely highlighting their hypothesized protective role for neurons in neurodegenerative diseases, potentially through increased degradation of the aggregate-prone peptides^57–59^. Neurons also demonstrated increased incorporation of D_2_O and d-glucose (**Fig. 5b**, MAP2, blue), reflecting possible upregulated *de novo* metabolic activity to counter aggregation toxicity, as most aggregates form within neurons. These findings emphasize the intricate interplay between protein aggregation, protein and lipid metabolism, and cellular stress responses, with potential to shedding new light in understanding the metabolic aspects in neurodegeneration.

## Discussion

Our platform provides a powerful tool for non-destructive, high-resolution, and quantitative metabolic imaging in live neuronal co-cultures with mixed neuronal types that have diverse morphology. By integrating label-free and bioorthogonal stimulated Raman scattering (SRS) microscopy with deep learning analysis, we enabled visualization of cell-type-specific metabolic activities with minimal perturbation. The platform’s capacity for real-time metabolic profiling, along with the flexibility to incorporate multiple deuterium-labeled metabolites, offers unprecedented insights into cellular metabolism and interaction dynamics. Our framework bridges the gap between 2D imaging approaches and exhaustive full-stack acquisition. By predicting high-fidelity 3D volumes from sparse data, it provides a practical sweet spot for enabling comprehensive volumetric analysis while minimizing photodamage and acquisition time.

Our approach integrates high-speed SRS imaging with a tandem RCNN and U-Net architecture for image reconstruction and cell-type-specific segmentation. This combined strategy significantly reduces data acquisition time while maintaining spatial resolution and cell viability. The model’s ability to interpolate sparse image stacks and perform accurate semantic segmentation underscores its potential for high-throughput metabolic screening in both basic research and drug discovery. Although direct training on live-cell images paired with corresponding antibody-stained images may seem ideal, the staining process can introduce re-embedding errors and artifacts. To mitigate these issues, we first train the model using fixed antibody-stained images paired with corresponding fixed-cell SRS images. Subsequently, we fine-tune the model using live-cell SRS data paired with fixed antibody-stained images. This two-stage training strategy enhances the model’s performance and extends its applicability to dynamic, living systems.

SRS imaging offers distinct advantages over conventional fluorescence microscopy and other label-free imaging techniques for metabolic profiling. Unlike fluorescence-based methods that often require genetically encoded sensors or fluorophore conjugates, SRS enables the probing of metabolic activities with minimal labeling. The use of deuterium-labeled molecules enhances this specificity, facilitating the visualization of metabolic pathways such as lipid synthesis, glucose metabolism, and protein turnover. The small molecular size of deuterium-labeled molecules compared to fluorescence probes minimizes concerns about metabolic perturbation, allowing for more accurate metabolic profiling.

A major challenge for metabolic imaging of neuronal co-cultures is the ability to distinguish individual cells, as neuronal cell shapes are not well-defined and can become tangled together. To address this, improving the deep learning model for enhanced morphological feature extraction, combined with more diverse training datasets, could improve cell boundary distinction and segmentation accuracy, enabling more precise and comprehensive analysis.

## Methods

### Reagents and materials

Nuclease-free water, bovine serum albumin (BSA), and anti-OLIG2 antibody (NC1331218) were acquired from Thermo Fisher. Paraformaldehyde (PFA, 16% in water) was purchased from Electron Microscopy Sciences. Anti-GFAP antibody (G3893) was purchased from Sigma-Aldrich. Anti-NG2 antibody (ab275024) and anti-MAP2 (ab5392) were purchased from abcam. Secondary antibodies: goat anti-rat IgG, Alexa Fluor 568 (Invitrogen, A-11077); goat anti-mouse IgG, Alexa Fluor 647 (Invitrogen, A-21236); goat anti-rabbit IgG, Alexa Fluor 488 (Invitrogen, A-11034); goat anti-chicken IgY, Alexa Fluor 647 (Invitrogen, A-21449).

### Primary neuron culture

Primary rat hippocampal neurons were isolated from neonatal Sprague–Dawley rat (CD (Sprague–Dawley) IGS rat, Charles River) pups with a protocol (IA22-1835) approved by Caltech’s Institutional Animal Care and Use Committee (IACUC). The brains were dissected from the skull and placed into a 10-cm Petri dish with ice-chilled Hanks’ balanced salt solution (Gibco). The hippocampus was isolated from the brains under a dissection scope, cut into small pieces (∼0.5 mm), and incubated with 5 ml of Trypsin-EDTA (0.25%, Gibco) at 37 °C with 5% CO_2_ for 15 min. The Trypsin-EDTA liquid was aspirated and replaced with 2 ml of DMEM containing 10% FBS to stop the digestion. The tissue fragments were moved into 2 ml of neuronal culture medium (Neurobasal A medium, B-27 supplement, 2 mM GlutaMAX supplement, Thermos Fisher, and 1× penicillin-streptomycin) and dispersed by repeated pipetting several times. The supernatant was collected and further diluted by neuronal culture medium to a final cell density of 9 × 104 cells ml^−1^. A 0.7-ml volume of cell suspension was added to each well of a 24-well plate on coated 12-mm circular cover glass. For pre-coating, sterile 12-mm circular cover glass was incubated with 100 μg/ml poly-d-lysine (Sigma) solution at 37 °C with 5% CO_2_ for 24 h in a 24-well plate. The 12-mm circular cover glass was washed twice with ddH_2_O and incubated with 10 μg/ml laminin mouse protein (Gibco) solution at 37 °C with 5% CO_2_ overnight. Thereafter, the 12-mm circular cover glass was washed twice with ddH_2_O and allowed to dry at room temperature inside a biosafety cabinet. Half of the neuron culture medium was replaced with fresh medium every 4 days. All the medium changes were done after pre-heating to 37°C.

### AAV packaging and AAV transduction

We used the pAAV backbone for building plasmid constructs. The inserted gene was a mutant HTT gene encoding the exon 1 of mHtt protein. The model mHtt protein contains an N-terminal 17 a.a. fragments (MATLEKLMKAFESLKSF), the polyQ tract (97Q), and the proline-rich region PPPPPPPPPPPQLPQPPPQAQPLLPQP QPPPPPPPPPPGP). For mHtt-EGFP protein, there is a linker (AVAEEPLHRPGSSPVAT) between mHtt and EGFP (MVSKGEELFTGVVPILVELDGDVNGHKFSVSGEGEGDATYGKLTLKFIC TTGKLPVPWPTLVTTLTYGVQCFSRYPDHMKQHDFFKSAMPEGYVQERTIFF KDDGNYKTRAEVKFEGDTLVNRIELKGIDFKEDGNILGHKLEYNYNSHNVYI MADKQKNGIKVNFKIRHNIEDGSVQLADHYQQNTPIGDGPVLLPDNHYLSTQ SALSKDPNEKRDHMVLLEFVTAAGITLGMDELYK*). The expression of mHtt protein was initiated by a general promoter, CMV (cytomegalovirus) promoter. The plasmids were sent to the CLOVER center at Caltech to produce AAV-DJ vectors. The typical titer is around 1E+13 genome copies (GCs)/mL with a normal yield of AAV vectors. AAV vectors were added to neurons at DIV 6-8 for about 5E9 GCs per well for the incubation of about 24 hours.

### Sample preparation and medium change

The sample preparation involved replacing the standard culture medium with a custom medium containing deuterium-labeled metabolites. D-OA and d-PA are added into the culture medium with a concentration of 20 μM. To enable the efficient incorporation, d_33_-oleic acid (Cambridge Isotope Laboratories, DLM-1891) and d_31_-Palmitic acid (Cambridge Isotope Laboratories, DLM-215) are coupled to bovine serum albumin (Sigma, A9418) in 2:1 molar ratio to prepare a 2 mM stock solution. The medium for d-AA, d-glucose, and D_2_O conditions, the medium was based on the Neurobasal A formula, with specific components replaced by deuterated analogs. The D-AA medium contained the following deuterated amino acids: D_7_-arginine·HCl (Cambridge Isotope Laboratories, DLM-541), D_8_-L-valine (Cambridge Isotope Laboratories, DLM-488-MPT), D_10_-L-isoleucine (Cambridge Isotope Laboratories, DLM-141), D_8_-L-tryptophan (Cambridge Isotope Laboratories, DLM-6903), D_9_-L, lysine·2HCl (Cambridge Isotope Laboratories, DLM-570), D_10_-L-leucine (Cambridge Isotope Laboratories, DLM-567), D_8_-L-methionine (Cambridge Isotope Laboratories, DNLM-7179), D_8_-L-phenylalanine (Cambridge Isotope Laboratories, DLM-372), D_5_-L-histidine·HCl·H2O (Cambridge Isotope Laboratories, DNLM-7366), D_3_-L-serine (Cambridge Isotope Laboratories, DLM-582), D_5_-L-threonine (Cambridge Isotope Laboratories, DLM-1693), D_4_-L-alanine (Cambridge Isotope Laboratories, DNLM-7178), D_7_-L-tyrosine (Cambridge Isotope Laboratories, DLM-589), and D_4_-L-cysteine (Cambridge Isotope Laboratories, DLM-9812). D_7_-glucose (Cambridge Isotope Laboratories, DLM-2062-1) was included with a concentration of 4.5 mg/ml and D_2_O was included with 30% of the water. For optimal incorporation, the medium with d-glucose, and D_2_O is added 5 days before imaging, the medium with 20 μM d-OA and d-PA is added 3 days before imaging, and the medium with d-AA is added 1 day before imaging. All the medium changes were done after pre-heating to 37°C.

### Immunostaining

The tissue-gel hybrid was transferred to 2 mL of PBST solution (0.1% Triton X-100 in PBS) and incubated with primary antibodies at a 1:100–1:200 dilution at 37 °C for 24 hours, followed by washing three times with PBST at 37 °C for 1–2 hours each. The samples were then incubated with secondary antibodies at a 1:200 dilution in PBST at 37 °C for 18–24 hours, followed by washing three times with PBST at 37 °C for 1–2 hours each before imaging.

### Fluorescence imaging

The fluorescence images of the immunolabeled samples were obtained with a Olympus Fluoview system. Images are captured with 4 μs pixel dwell time by a 25×, 1.05 NA water-immersion objective. Single-photon confocal Laser scanning imaging was performed with 488, 561, and 640 nm dye lasers (Coherent OBIS).

### SRS microscopy

The picoEmerald laser system (Applied Physics & Electronics) generated both the pump and Stokes beams used in this setup. The tunable pump beam had a wavelength range of 770 nm to 990 nm and a spectral bandwidth of approximately 7 cm^−1^, while the Stokes beam had a fixed wavelength of 1031.2 nm and a spectral bandwidth of 10 cm^−1^. These beams operated at an 80 MHz repetition rate, with the Stokes beam being modulated at 20 MHz via an integrated electro-optic modulator (EOM). Within the picoEmerald system, the pump and Stokes beams were synchronized both spatially and temporally before being directed into an Olympus FV3000 microscope. The Raman loss signal from the pump beam was demodulated using a lock-in amplifier (SR844 from Stanford Research Systems) at the modulation frequency. The in-phase output signal was fed back into the Olympus IO interface box (FV30-ANALOG) of the microscope. Image acquisition speed was set at 40 μs pixel dwell time and a 30 μs time constant. Images were captured in a 16-bit grey scale using Olympus Fluoview 3000 software.

For imaging, a 25× water immersion objective lens (XLPLN25XWMP, 1.05 NA, Olympus) focused the beams onto the sample. Transmitted light was collected using an oil immersion condenser lens (1.4 NA, Olympus). A bandpass filter (893/209 BrightLine, 25 mm, Semrock) was employed to block the Stokes beam, allowing only the pump beam to reach a 10 mm × 10 mm Si photodiode (S3590-09, Hamamatsu). To enhance the photodiode’s saturation threshold and reduce its response time, it was reverse-biased at 64 V. The output current was filtered through a 19.2–23.6 MHz bandpass filter (BBP-21.4+, Mini-Circuits) and terminated with a 50 Ω load.

### Image preprocessing

SRS images underwent lock-in background subtraction. Fluorescence images were segmented using CellProfiler with Otsu’s method to generate masks for each channel, which were then combined to create ground-truth images. Image alignment was performed using a custom ImageJ macro with a rigid body transform. The xy alignment was based on MultiStackReg, using the combined intensity of three fluorescence channels to match with the SRS images. Z-axis alignment was performed by identifying and matching the plane with the highest intensity. Finally, images were cropped into 488 × 488 patches to fit the model architecture.

### U-Net Deep Learning Model Implementation

The deep learning framework was implemented using pytorch, configured to operate on 3D high resolution datasets with nnUnet framework^60^. Deep supervision is applied, with auxiliary losses incorporated in the decoder for all but the two lowest resolutions. Data augmentation methods including rotations, scaling, Gaussian noise, Gaussian blur, brightness and contrast adjustments, low-resolution simulation, gamma correction, and mirroring are applied.

The loss function is a combination of cross-entropy and Dice loss. For deep supervision, downsampled ground truth segmentation masks are used at each resolution level. The total loss is computed as a weighted sum across all resolutions, where the weights decrease by half at each lower resolution level and are normalized to sum to 1.

Each U-Net follows a consistent configuration of two blocks per resolution step in both the encoder and decoder. Each block consists of a convolution, followed by instance normalization and a leaky ReLU activation. Downsampling is performed using strided convolutions, while upsampling uses transposed convolutions. To balance performance and memory efficiency, the initial number of feature maps is set to 32, doubling at each downsampling step and halving at each upsampling step. The maximum number of feature maps is capped at 320 to limit model size.

Each network is trained for at least 1,000 epochs, where one epoch consists of 250 mini-batches. Network weights are optimized using stochastic gradient descent with Nesterov momentum (µ = 0.99) and an initial learning rate of 0.01. Mini-batches are sampled randomly from training cases, with oversampling implemented to mitigate class imbalances. In each mini-batch, 66.7% of patches are selected randomly, while 33.3% are guaranteed to contain foreground classes. A minimum of one foreground patch per batch is enforced when the batch size is two.

### RCNN Deep Learning Model Implementation

The deep learning framework consists of a generator network and a discriminator network based on a convolutional recurrent network^39,61^. The network was implemented using TensorFlow and Keras with mixed precision computation. The generator and discriminator networks were constructed using convolutional and recurrent layers to process image sequences. The generator model is based on a multi-scale convolutional-recurrent network. The architecture consists of an initial convolutional block followed by a downsampling path, a bottleneck block, and an upsampling path.

The input processing stage reshapes the input tensor and passes it through two convolutional layers with ReLU activation to extract initial features. The downsampling path comprises multiple DownBlocks, each consisting of convolutional layers, batch normalization, and leaky ReLU activation. Max pooling layers reduce the spatial resolution while retaining essential features. The bottleneck layer, composed of a DownBlock and a recurrent convolutional block (RCBlock), processes the lowest-resolution feature maps. The upsampling path includes multiple UpBlocks, each incorporating transpose convolution layers, batch normalization, and leaky ReLU activation. Skip connections are used to concatenate features from corresponding downsampling layers. Finally, the output processing stage generates the final output using two Conv2D layers, where the last layer applies a linear activation function.

The generator’s loss function is a weighted sum of Mean Absolute Error (MAE), Structural Similarity Index (SSIM), and, Total Variation Loss (TV Loss). Mean Absolute Error (MAE) quantifies pixel-wise differences between the generated and target images. The Structural Similarity Index (SSIM) ensures the generated images maintain perceptual similarity to the target. Additionally, Total Variation Loss (TV Loss) encourages spatial smoothness in the generated images. The discriminator loss is measured using mean squared error (MSE) between the generated and real data.

The discriminator model is a convolutional neural network designed to differentiate between real and generated images. Its architecture includes an initial convolutional layer, which applies a Conv2D layer with ReLU activation to extract features. This is followed by normal blocks, a series of convolutional layers that progressively increase the feature depth while reducing spatial resolution. A global average pooling layer reduces feature maps to a single vector, which is then processed by fully connected layers—two dense layers, with the final one using a sigmoid activation function to produce a probability score.

The training process follows an adversarial learning framework where the generator and discriminator are updated iteratively. Loss gradients are computed using TensorFlow’s automatic differentiation, and the Adam optimizer is employed for weight updates. The model was trained using mini-batch stochastic gradient descent. Inference was performed using the trained generator to produce synthesized images.

### Image analysis

CD images were segmented using the predicted mask, with non-segmented areas classified as background. For each dataset, background-subtracted values were calculated by subtracting the mean background intensity from the corresponding dataset. To ensure data quality, datasets were filtered based on a minimum pixel count of 400 (∼100 µm²) per category, preventing disproportionate representation by a small portion of a cell or a single cell within the dataset, and to ensure a mixed co-culture environment. Statistical comparisons between experimental conditions were performed using analysis of variance (ANOVA) to assess significance. Droplet counting was conducted using CellProfiler with EnheanceOrSuppressFeatures module (enhance speckles of feature size 2), followed by IdentifyPrimaryObjects thresholded with Robust Background. The center of each droplet was recorded and categorized based on the predicted label of the corresponding pixel^62^.

## Author contributions

L.W. supervised the project. L.-E.L. and L.W. conceptualized and designed the research. L.-E.L. conducted the experiments and analyzed the data. X.B. cultured the cell samples and introduced perturbations. A.C. optimized the imaging equipment. H.W. optimized image acquisition methods. L.-E.L. and L.W. wrote the manuscript, with contributions and revisions from all authors.

## Acknowledgment

We acknowledge the Caltech Institutional Animal Care and Use Committee (IACUC), and the Office of Laboratory Animal Resources (OLAR) for research resources. This work is supported by a CZI dynamic imaging grant and a Vallee Scholars award. L.-E.L. acknowledges the support received from the J Yang & Family Foundation Graduate Fellowships at Caltech. X.B. acknowledges the support received from the Biotechnology Leadership Pre-Doctoral Training Program (BLP) in the Donna and Benjamin M. Rosen Bioengineering Center and the Barbara J. Burger Graduate Fellowship at Caltech. A. Colazo acknowledges the support received from National Science Foundation Graduate Research Fellowship under Grant No. 2139433. L.W. is a Heritage Principal Investigator supported by the Heritage Medical Research Institute.

