## Supplementary information for "Deep Learning-Augmented Stimulated Raman Imaging for Cell-Type-Specific Metabolic Profiling in Live Neuronal Co-Cultures"

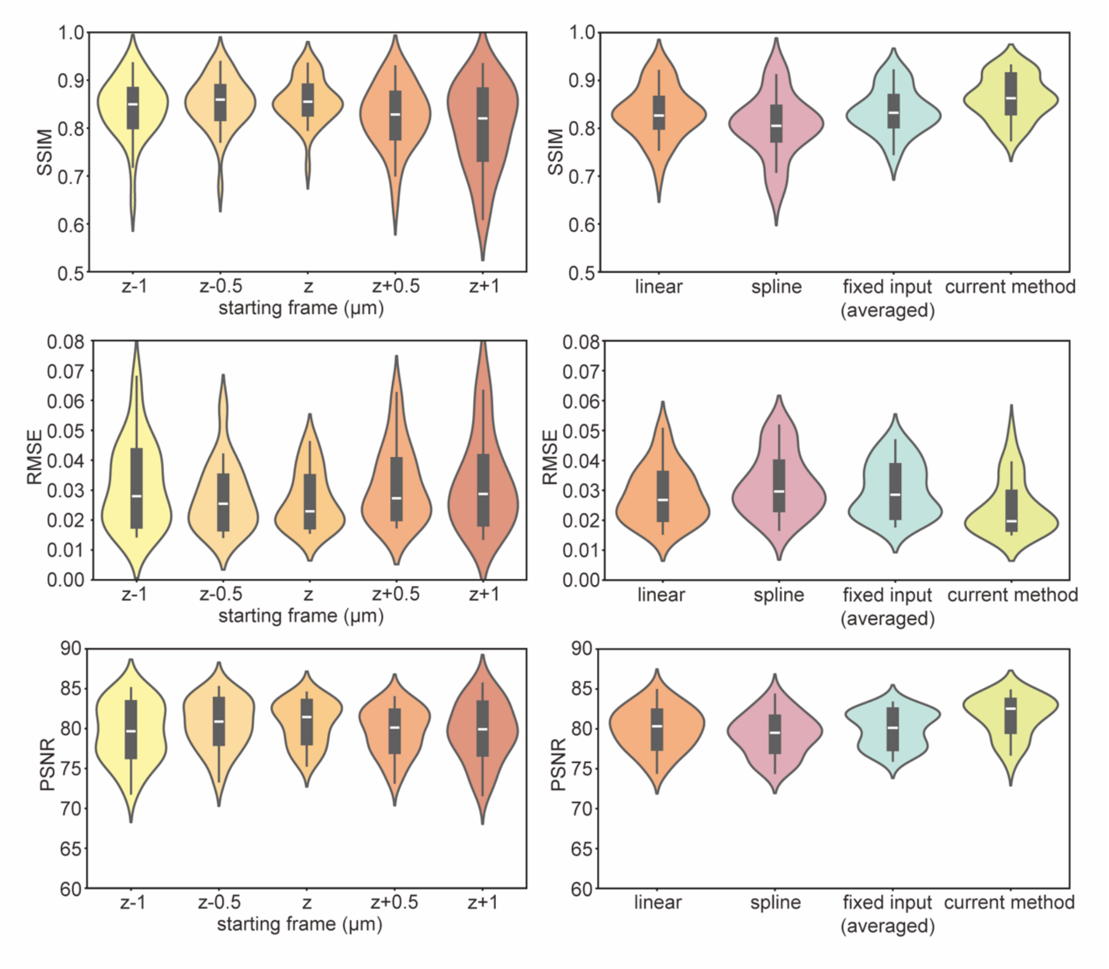


**Figure S1. Comparison of model performance metrics, including structural similarity index measure (SSIM), root mean square error (RMSE), and peak signal-to-noise ratio (PSNR) of each model.** (Left) Performance matrices of models trained using different fixed starting frames relative to the cell body, based on data from a complete 0.5 μm interval stack. (Right) Performance matrices of stacks generated by linear interpolation ("linear"), spline interpolation ("spline"), the average performance of five models trained on specific fixed starting frames ("fixed input, averaged"), and our current model ("current method") trained on mixed-starting-frame stacks.

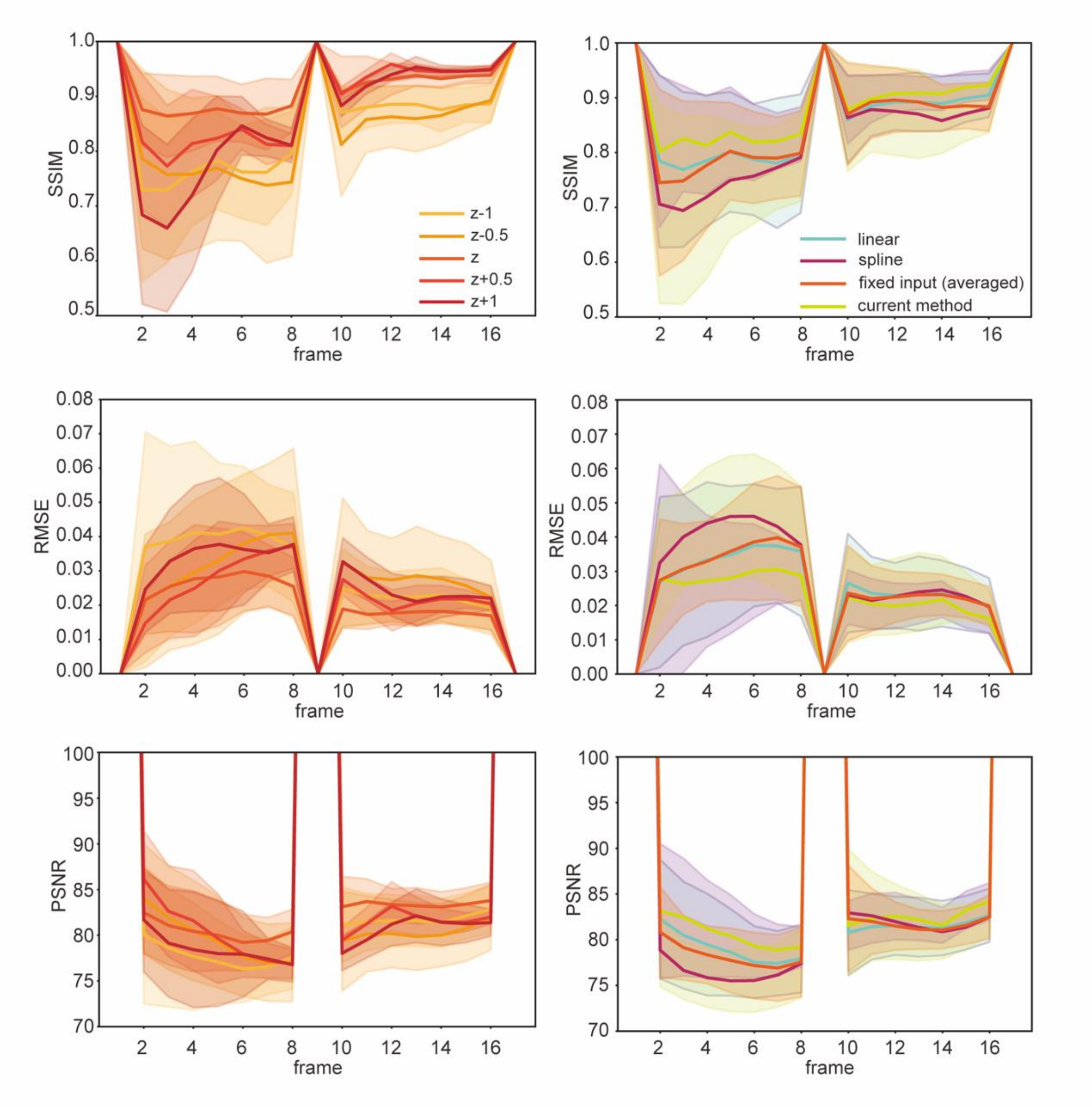


**Figure S2. Comparison of model performance metrics, including SSIM, RMSE, and PSNR of each frame.** (Left) Performance matrices of models trained using different fixed starting frames relative to the cell body, based on data from a complete 0.5 μm interval stack. (Right) Performance matrices of stacks generated by linear interpolation ("linear"), spline interpolation ("spline"), the average performance of five models trained on specific fixed starting frames ("fixed input, averaged"), and our current model ("current method") trained on mixed-starting-frame stacks. Frames 1, 9 and 17 are input images.

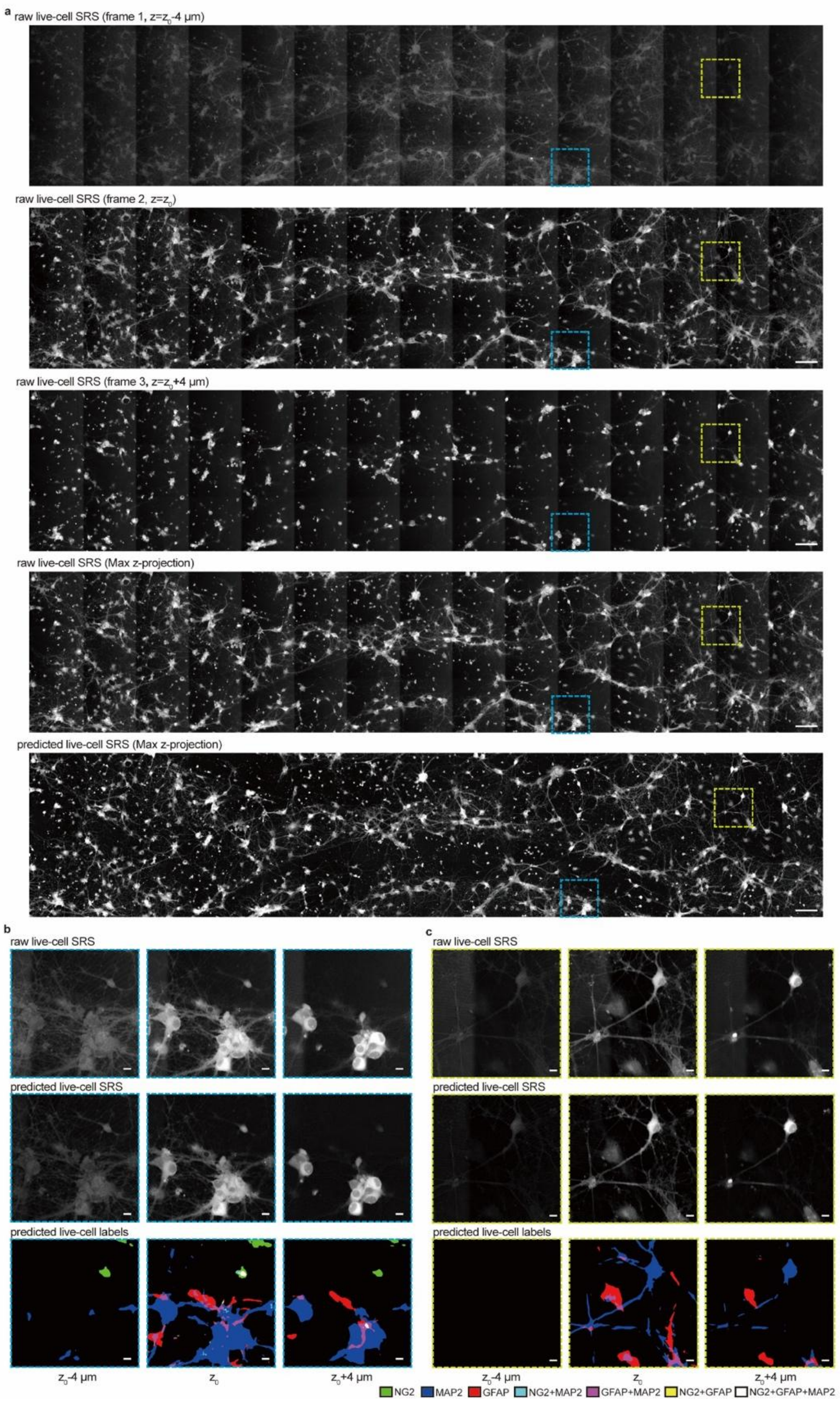


**Figure S3. Representative images with model prediction**. (a) Overview of 3 raw SRS frames from a mixed neuronal culture, along with the maximum z-projection of the 3 frames, and the maximum z-projection of the predicted 17-frame SRS stack. (b-c) Raw SRS frames, corresponding predicted SRS frames, and corresponding predicted labels of the highlighted regions in (a, blue dashed (b) and yellow dashed (c) boxes), showing overlapped cells (b) and isolated cells (c). Scale bars: (a), 100 µm; (b-c), 10µm.

**Table S1. Performance metrics of the model for neurons (MAP2) prediction, trained with both full and under sampled data.**

| Sampling methods | Accuracy | Dice | Precision | Recall |
|---|---|---|---|---|
| Whole stack | 0.9790±0.0175 | 0.7379±0.1025 | 0.7752±0.0992 | 0.7147±0.0801 |
| Under sampled | 0.9724±0.0207 | 0.6203±0.1001 | 0.7085±0.1303 | 0.5696±0.0904 |
| RCNN-restored | 0.9774±0.0194 | 0.7104±0.0754 | 0.7734±0.0511 | 0.6611±0.0999 |

**Table S2. Performance metrics of the model for astrocytes (GFAP) prediction, trained with both full and under sampled data.**

| Sampling methods | Accuracy | Dice | Precision | Recall |
|---|---|---|---|---|
| Whole stack | 0.9754±0.0152 | 0.7318±0.0804 | 0.7512±0.0840 | 0.7148±0.0821 |
| Under sampled | 0.9695±0.0172 | 0.6564±0.0820 | 0.6964±0.1103 | 0.6283±0.0786 |
| RCNN-restored | 0.9737±0.0166 | 0.7036±0.0869 | 0.7348±0.0832 | 0.6775±0.0947 |

**Table S3. Performance metrics of the model for oligodendrocytes (OLIG2) prediction, trained with both full and under sampled data.**

| Sampling methods | Accuracy | Dice | Precision | Recall |
|---|---|---|---|---|
| Whole stack | 0.9987±0.0014 | 0.8819±0.1030 | 0.8791±0.1035 | 0.8857±0.1061 |
| Under sampled | 0.9977±0.0027 | 0.7609±0.1195 | 0.8777±0.1190 | 0.6838±0.1430 |
| RCNN-restored | 0.9988±0.0014 | 0.8679±0.1462 | 0.8718±0.1424 | 0.8774±0.1231 |

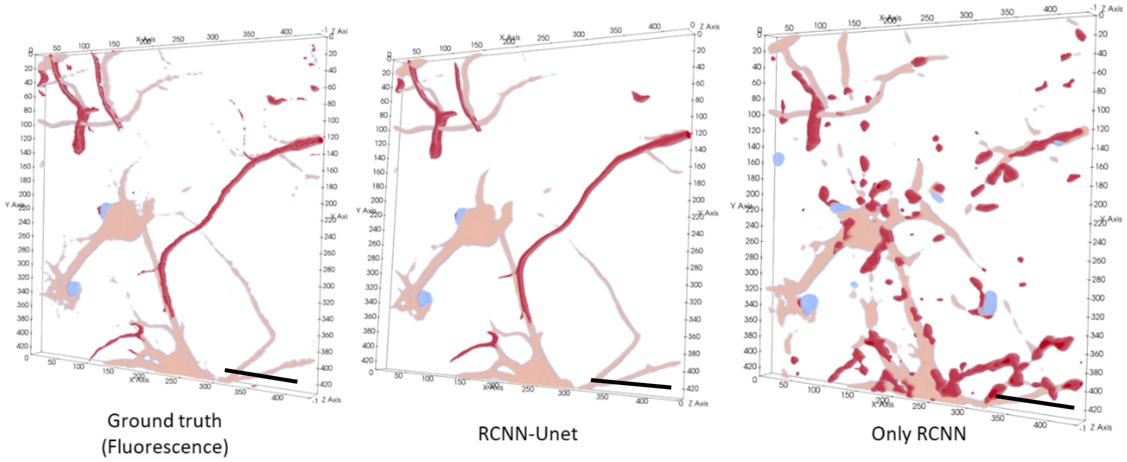


**Figure S4. Direct application of RCNN to predict the entire fluorescence stack using a three-frame SRS input results in poor model performance.** Even with extended training, the model remained constrained to predicting fractured patterns in the labels (Only RCNN compared to Ground truth). Scale bars: 100 µm.

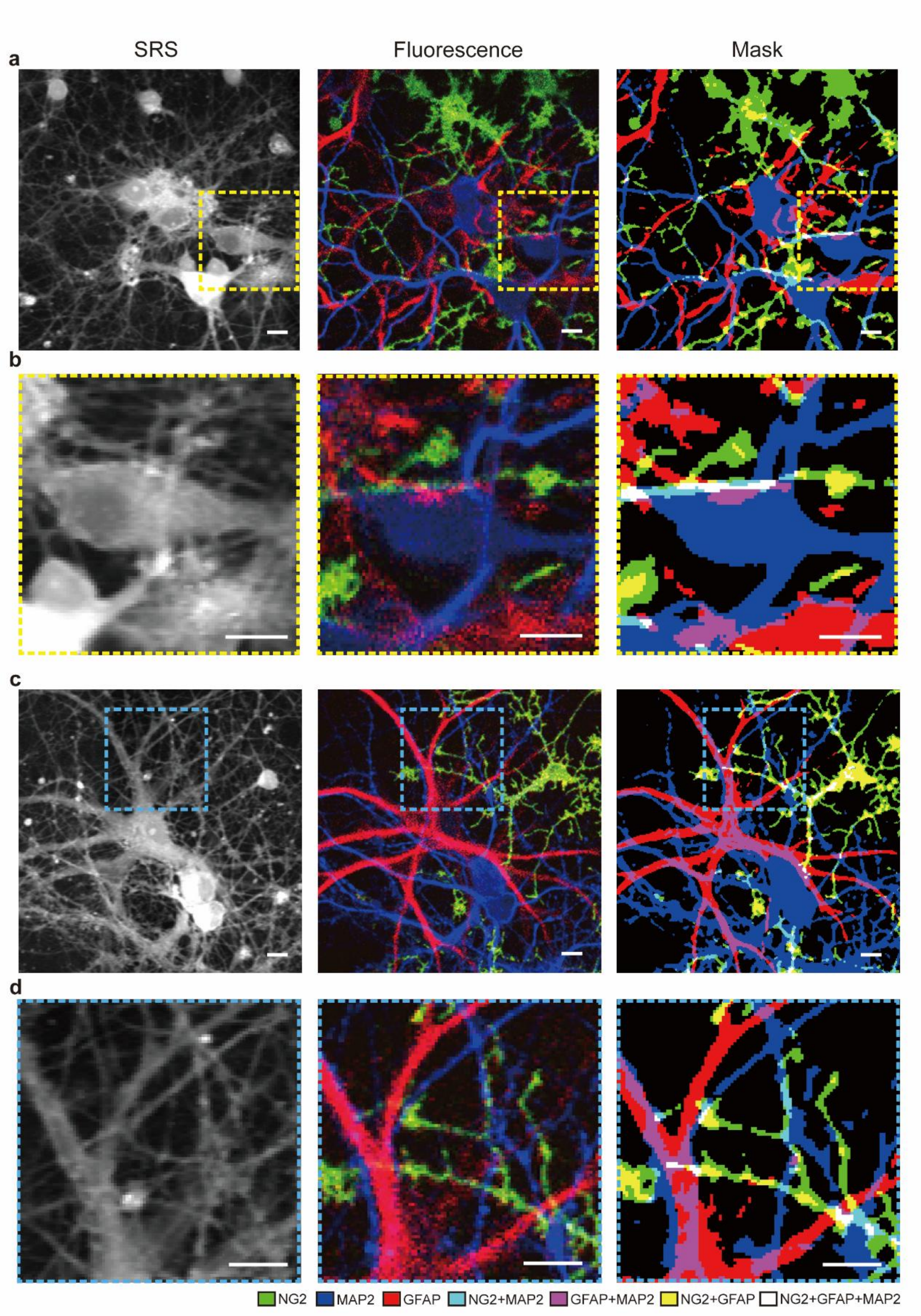


**Figure S5. SRS images (left), fluorescence images (middle), and predicted masks (right) from regions containing overlapping pixel labels for oligodendrocytes (NG2), astrocytes (GFAP), and neurons (MAP2).** Panels (a) and (c) show representative overlapping regions, while panels (b) and (d) show the corresponding enlarged views. Scale bars: 10 μm.

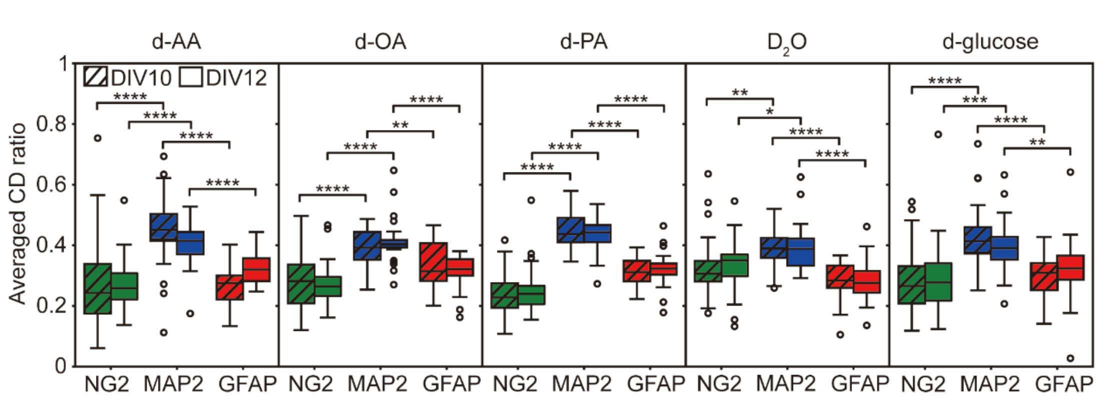


**Figure S6. Statistical comparison of metabolic activities across neuronal cell types for data shown in Fig. 2a.** Relative CD uptake ratio among predicted oligodendrocytes (NG2), neurons (MAP2), and astrocytes (GFAP) between imaging at DIV10 and DIV12 for D-AA (both 1-day incubation), D-OA, D-PA (both after 3-day incubation), $D_2O$, and D-glucose (both after 5-day incubation). Neurons showed higher relative metabolic activities for all metabolites. The sum of CD ratios from all three cell types is equal to 1. *, $P \leq 0.05$; **, $P \leq 0.01$; ***, $P \leq 0.001$; ****, $P \leq 0.0001$.

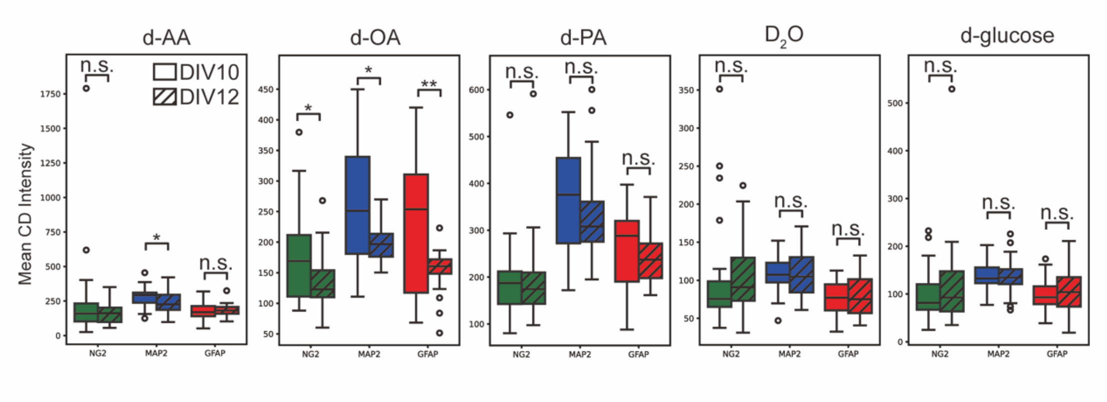


**Figure S7. Comparison of CD intensity for assessing the absolute metabolic uptake and incorporation of deuterated metabolites imaged on DIV10 and DIV12.** The medium was replaced with deuterated molecule-containing medium 1 day before imaging for d-AAs, 3 days for d-OA and d-PA, and 5 days for $D_2O$ and d-glucose. n.s., $P > 0.05$; *, $P \leq 0.05$; **, $P \leq 0.01$.

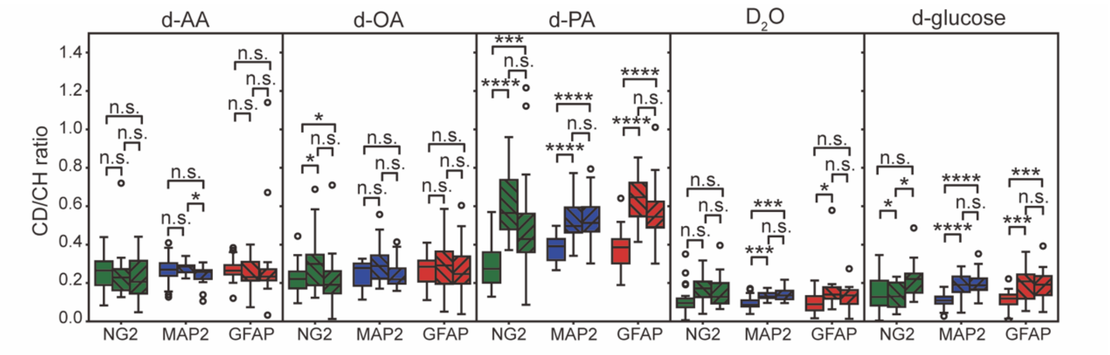


**Figure S8. Comparison of CD/CH ratio across three groups: non-transduced cultures (Regular), AAV-transduced cultures with protein aggregation (w/ agg.), and AAV-transduced neuronal cells without protein aggregation (w/o agg.).** n.s., $P > 0.05$; *, $P \leq 0.05$; **, $P \leq 0.01$; ***, $P \leq 0.001$; ****, $P \leq 0.0001$.
